# Infrared absorptions as indicators for *Pseudomonas aeruginosa* and *Acinetobacter baumannii*

**DOI:** 10.1101/2020.03.25.007591

**Authors:** Masato Yamamoto, Satoru Arata, Kunihiko Fukuchi, Hidehiko Honda, Hirokazu Kobayashi, Masahiro Inagaki

**Author notes:** Corresponding author, (MY).

## Abstract

There is clinical demand for simple and contact-free diagnosis techniques in medical practice. This study shows that the infrared absorptions of volatile metabolites can be used to distinguish between the air around *Pseudomonas aeruginosa, Acinetobacter baumannii,* other bacteria, and normal room air. Gas samples were collected from the air surrounding single and mixed laboratory cultures, and preliminary data on human breath samples are also presented. The infrared spectra of a variety of gasses are measured with a high resolution of 0.5 cm^-1^ to obtain information about the wavenumber position of the key bands. Hence, it is not necessary to specify the molecular species of biomarkers. This work also shows that discrimination rates can be improved by observing additional infrared absorptions caused by different characteristic volatile molecules. The significance of this work is that the specific wavenumber positions of the key bands that allow the application of infrared lasers are provided. As a result, it is considered that *Pseudomonas aeruginosa* and *Acinetobacter baumannii* can be monitored more sensitively and easily. With further research and development, this simple approach could be used in future applications to identify infections in healthcare settings.

## Introduction

Biomarkers are characteristic molecules produced by biological processes. The biomarkers found in various gas samples have been examined by using gas chromatography–mass spectrometry (GCMS), and they are of interest for non-invasive diagnosis and healthcare applications [1–5]. In addition, representative small molecules such as CO, CO_2_, NO, and NO_2_ have been the target of high-sensitivity infrared laser spectroscopy measurements [6–9]. Clinical applications of these techniques have been proposed [10–12].

To apply an infrared laser, it is necessary to adjust the oscillation wavenumber (the reciprocal of the wavelength) to a specific value. A molecule produces many peaks, but the wavenumber indicates the position of one strong peak on the horizontal axis of the infrared spectrum. Infrared lasers can be used to detect changes in the concentration (partial pressure) of some target volatile biomarkers. In particular, it has been shown that changes in the concentration of ethane [13] and ethanol [14] can be detected with ultra-high sensitivity, in addition to the molecules CO, CO_2_, NO, and NO_2_.

High resolution infrared spectra can be obtained for a wide variety of gas samples. By analyzing the infrared absorption peaks it is possible to identify the key band, the infrared absorption peak that serves as the target index of a biomarker. There are two criteria for defining a key band. First, it must appear in a “window” region where there are no unrelated strong peaks; this means the intensity of the key band can be observed clearly. Second, the peak position of the wavenumber must be accurately determined in a range close to the linewidth of the infrared laser.

The intensity and wavenumber position of the key band correspond to quantitative information about the concentration (partial pressure) of key volatile metabolites (VMs) or volatile organic compounds (VOCs). This information is the index equivalent to surveillance markers and can be used to distinguish between bacterial species using the surrounding air.

In recent years, several reports [15–19] have classified bacteria based on infrared spectra measured using Fourier-transform infrared (FTIR) spectroscopy. In these studies, samples were prepared by treating cultured bacteria, and the analysis target was the bacteria itself, not the VMs released from the bacteria. The infrared spectra were measured with a general resolution of 4 cm^-1^ by directly irradiating bacteria with infrared light and transmitting or reflecting the light.

In this study, infrared spectra have been recorded for a wide variety of gas samples including gases obtained near eight types of bacteria and reference samples of indoor air. Each of these gas samples contains a wide variety of molecules, and each molecule (except for homonuclear diatomic molecules such as O_2_ and N_2_) produces many infrared absorption peaks. An infrared spectrum for one gas sample may include thousands of peaks in the range of 550–7000 cm^-1^. By analyzing the accumulation of the infrared spectra, the wavenumber positions of the key bands (the characteristic infrared absorption peaks) can be found. Therefore, the oscillation frequency (wavenumber) of the infrared laser can be fine-tuned in a limited area, which makes it possible to detect bacteria with high sensitivity without identifying the molecular species as the target index markers.

The characteristic molecules released into the atmosphere by some types of bacteria have also been investigated using GCMS [20]. In particular, methyl thiocyanate and benzonitrile have been identified as characteristic VOCs emitted by *Pseudomonas aeruginosa*. These molecules give strong infrared absorption peaks in the region of 2000–2300 cm^-1^. However, specifying the molecular structure of the key VOC does not automatically mean it can be found because the key band must appear in the “window” region so that the intensity can be read. The condition is confirmed only by comparison with many infrared spectra of other gas samples.

The next condition is whether the accuracy of the key-band position is in a range that facilitates the application of an infrared laser. Experimental results for a pure substance and the theoretical calculations for a molecule, such as normal vibration analysis for the most stable structure, may be used to predict candidates for the key bands. However, gas samples contain a mixture of substances including H_2_O, CO_2_, and other atmospheric components, which could affect the key-band position by binding to the key molecule. Furthermore, the infrared spectra may potentially reflect molecular aggregates, clusters, and small particles suspended in the atmosphere, which have a different structure than that suggested by the GCMS. A key molecule does not always give the key bands at the same position, intensity, and width as the prediction.

This study aims to identify the key bands that characterize the air around *Pseudomonas aeruginosa* and *Acinetobacter baumannii*. These are typical drug-resistant bacteria that are associated with nosocomial infections, so there is urgent need for simple and contact-free detection techniques that can be used in medical practice. The wavenumber positions of the key bands can be used to discriminate between gas samples without specifying the molecular species of the biomarkers.

Specifically, the wavenumber position of the key bands for VOCs or VMs emitted by the bacteria are recorded at a high resolution of 0.5 cm^-1^, which makes it possible to determine the type of bacteria without contact. Furthermore, this work aims to demonstrate that the high intensity infrared absorption peak that appears at a specific wavenumber position can be used as an indicator for nearby bacterial growth, even if the molecular structure of the characteristic VOC(s) is not specified.

The positions of the key bands are found from the experimental results (the accumulation of infrared spectra in the region of 550–7000 cm^-1^) for various mixed substances, as a result of the search and selection. Therefore, the key substance is not limited to a single molecule in the most stable structure. This differs from the previous studies because any suspended substances shown by the infrared absorption peaks are covered.

## Methods

### Preparation of gas samples

Gas samples were obtained from the air surrounding cultured bacteria. The gases were aspirated into gas bags (smart bag PA, smart bag 2F, Tedlar bag or ANALYTIC-BARRIER bag, GL Sciences) by the indirect sampling method using a dry pump. The gas bag was equipped with a standard sleeve (6-7 mm O.D.) connected to a silicone tube within 1 m. The other end of the silicone tube was placed within 5 cm of the object. Eight types of bacteria, including standard strains and hospital isolates, were cultured in sheep’s blood agar at 37 °C for 30 to 40 h. The details are summarized in Table 1. *M1, M2, M3*, and *M4* indicate mixed gas samples obtained from plural types of bacteria cultured in a nutrient agar at 37 °C. For comparison, normal room air, human breath, and air from the vicinity of the culture medium alone were also obtained and measured. The human breath was blown into a polyethylene zipper bag.

**Table 1.**
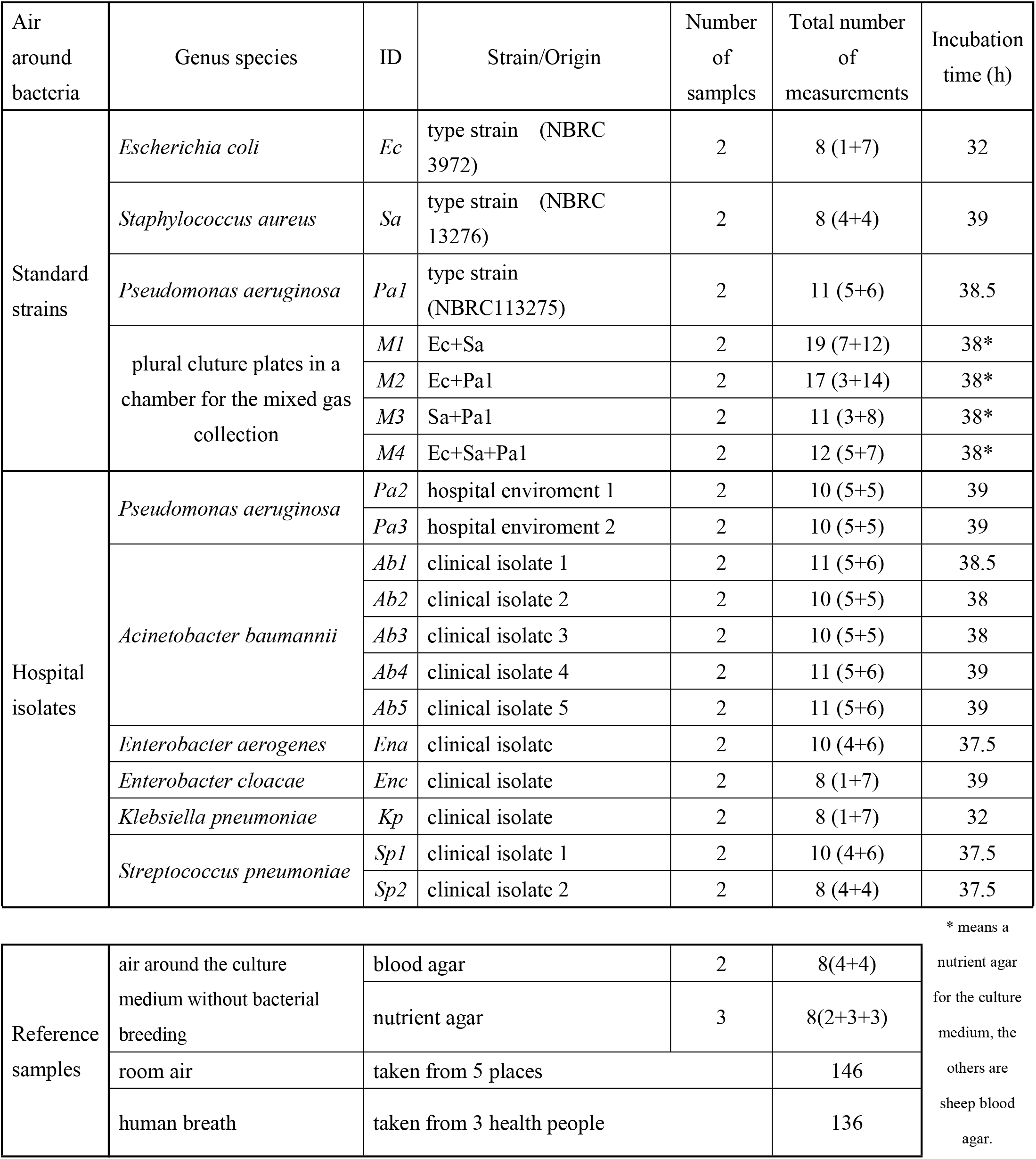
Details of the gas samples.

### Measurements

The gas in each bag was introduced into a gas cell (light path length 10 m) installed in an FTIR spectrometer (Bruker VERTEX 70). Overall, 365 infrared (IR) spectra for the various gas samples in Table 1 were measured in the range of 550–7500 cm^-1^ at atmospheric pressure with a resolution of 0.5 cm^-1^.

### Data analysis

Microsoft EXCEL was used for data processing and analysis. The details of the infrared absorption peaks that appeared in the IR spectra were obtained and the peak intensities were measured and compared. The peak intensity was calculated by subtracting the absorbance at the base point from that at the absorption peak. Peak and base pairs were found around the peak positions where the target group showed strong absorption. The mean and variance of the absorption intensity for the reference group were compared with the target group. Peak and base pairs with high discrimination rates were extracted for each, pair and high discrimination rates were achieved for *P. aeruginosa* and *A. baumannii.*

### Ethics statement

This study was approved by the research ethics committee of Showa University School of Medicine (Approval No. 371 and 2510).

## Results and discussion

### Pseudomonas aeruginosa (Pa)

Fig 1 shows the IR spectra at approximately 2213 cm^-1^ for the air surrounding the eight types of cultured bacteria and the normal room air. These nine spectra are offset and aligned. The vertical axis indicates absorbance and the absorption peak appears in the upward direction. In this region of the IR spectra, a strong peak can be observed at 2213.2 cm^-1^, which is characteristic of *Pa*.

**Fig 1.**
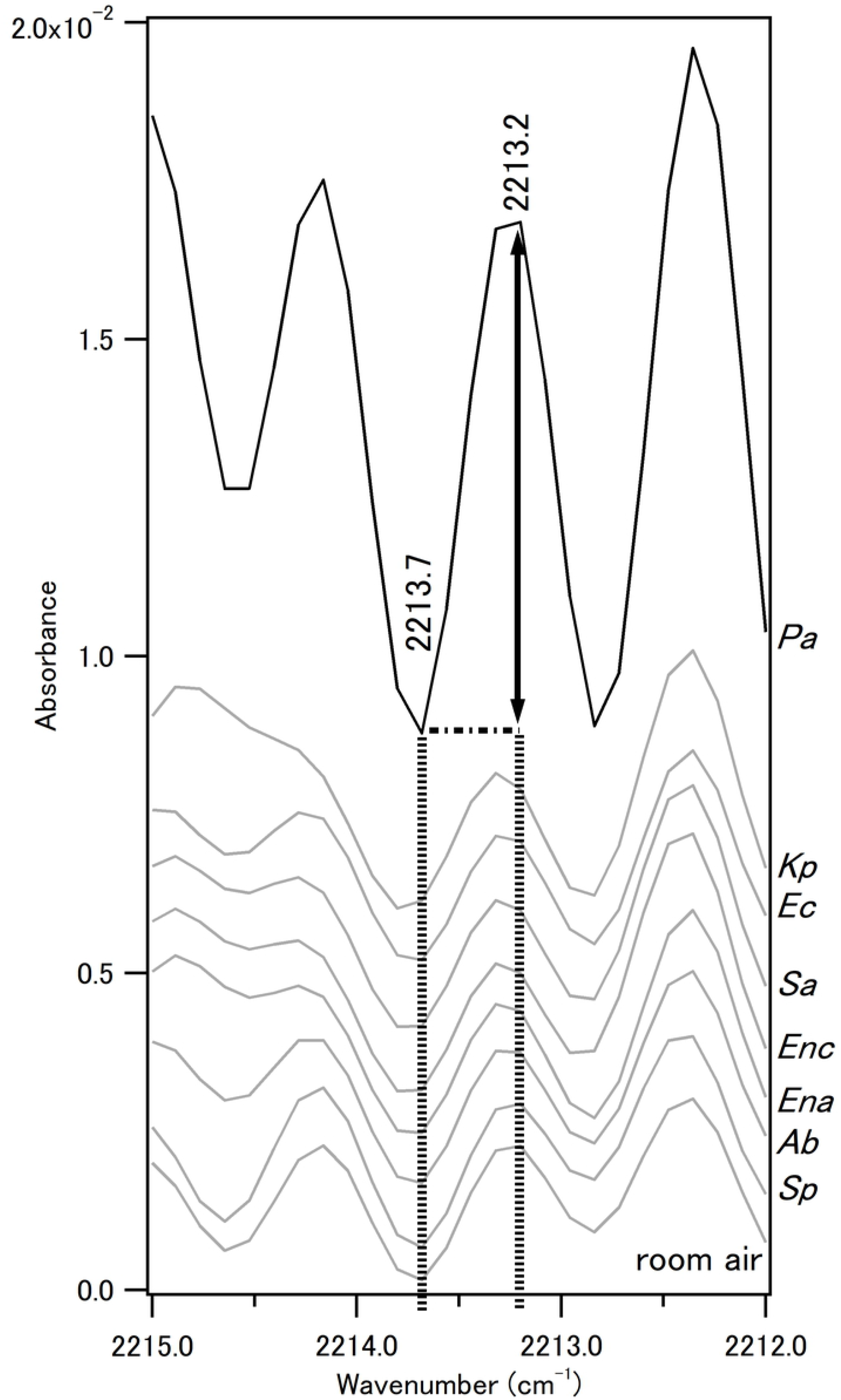
IR spectra at approximately 2213 cm^-1^ for the air surrounding 8 types of bacteria and room air. The 9 spectra are offset and aligned in the following order from the top: *Pseudomonas aeruginosa (Pa), Klebsiella pneumoniae (Kp), Escherichia coli (Ec), Staphylococcus aureus (Sa), Enterobacter cloacae (Enc), Enterobacter aerogenes (Ena), Acinetobacter baumannii (Ab), Streptococcus pneumoniae* (*Pa*), and room air.

As shown by the two dotted lines in Fig 1, two positions in the wavenumber (horizontal axis) were fixed to determine the relative intensity of the absorption peak (P) at 2213.2 cm^-1^ (P at 2213.2) compared to the value in absorbance of the base point (B) at 2213.7 cm^-1^ (B at 2213.7). As indicated by the two-way arrow in Fig 1, the difference was calculated by subtracting the absorption value at the base point from that at the peak. This value was used as the intensity of the absorption peak at 2213.2 cm^-1^ assigned to the VM from *Pa.* In all of the spectra, the two fixed positions for the assumed peak and base do not actually correspond to the wavenumbers at the local maximum and minimum, respectively.

The absorption peaks at 2213.2 cm^-1^ and 778.4 cm^-1^ were taken as the index of *Pa.* The absorption intensity of the peak at 778.4 cm^-1^ (P at 778.4) is the difference from the value for the base point at 778.6 cm^-1^ (B at 778.6).

Fig 2 shows the results for discrimination of *Pa* using two intensities as the *Pa* indexes. The subtracted value, P at 778.4 – B at 778.6, is shown on the horizontal *(x)* axis, and that of P at 2213.2 – B at 2213.7 is shown on the vertical (*y*) axis. The gas samples that included VMs from *Pa* (filled grey squares 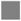) are grouped in the upper right region, which is delimited by two dotted lines (*x* > 0 and *y* > 0.003). All of the other cases are distinguished. The 365 points in Fig 2 correspond to the values obtained for the 365 spectra from the various gas samples shown in Table 1. Open circles 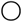 indicate the values obtained from human breath samples. The infrared spectra of human breath were measured under the same conditions as the air samples for cultured bacteria and they have been stored for analysis. The results will be published in later works.

**Fig 2.**
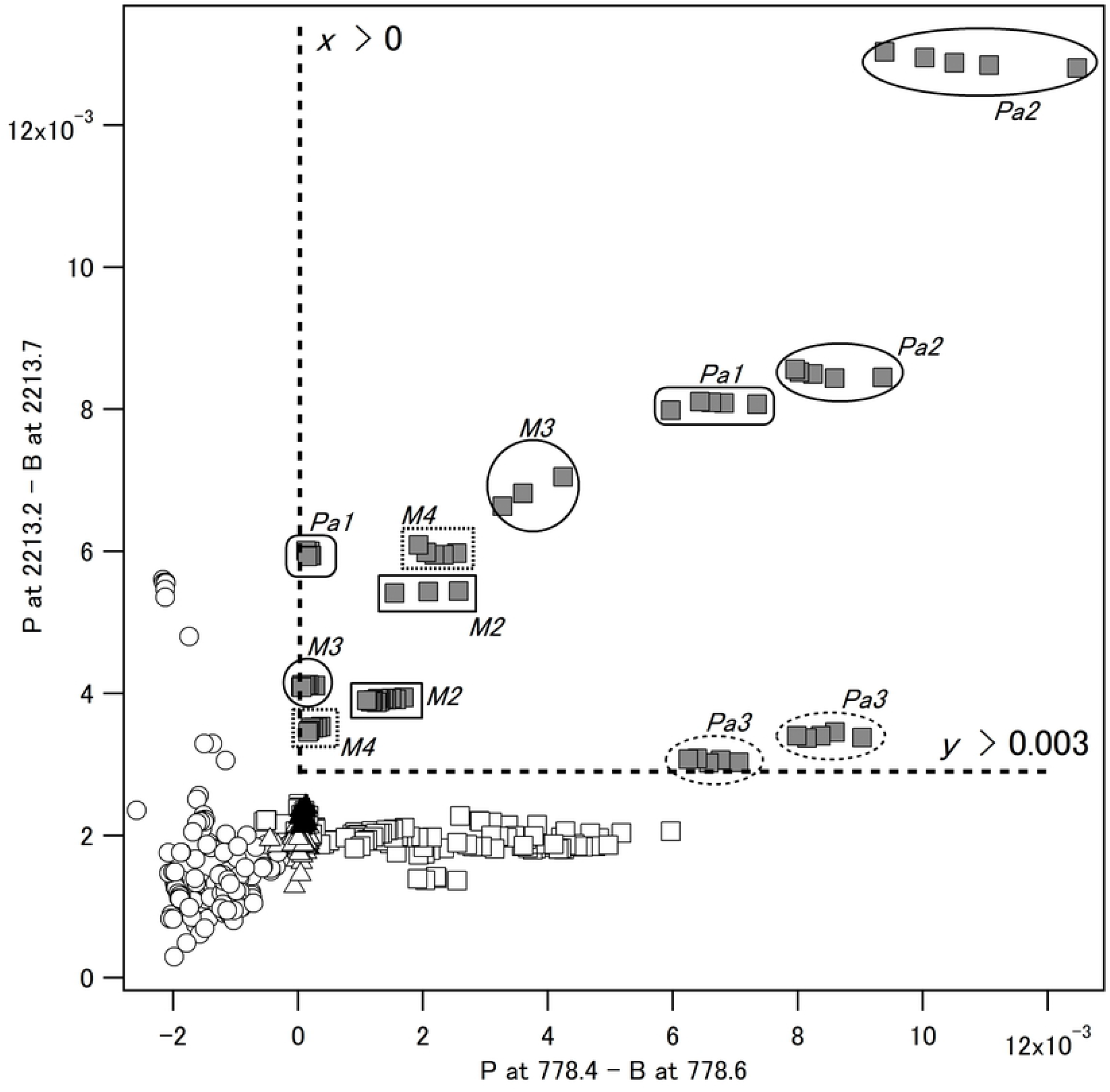
Discrimination of *Pseudomonas aeruginosa (Pa)* with two indexes, P at 778.4 - B at 778.6 and P at 2213.2 - B at 2213.7, shown on the horizontal (*x*) and vertical (*y*) axes, respectively. Grey squares 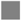, the values obtained from the gas samples that included VMs from *Pa;* open squares □, the other 7 types of bacteria; black triangles ▲, air from the vicinity of the culture medium alone; open triangles Δ, the normal room air; and open circles 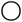, human breath. *M2, M3,* and *M4* indicate the data from the mixed gas samples that included VMs from *Pa* and the other type(s) of bacteria, as shown in Table 1.

The values on the *x* and *y* axes reflect the infrared absorption peaks of the VMs specific to *Pa*. According to Lambert-Beer’s law, the infrared absorption intensity indicated by absorbance is proportional to the concentration or partial pressure. The gas samples contain multiple unknown chemical species and the absorption peaks from the different components could overlap in the infrared spectrum. In this case, the absorbance intensity would not show a simple proportional relationship. The main VM(s) on the *x* axis is (are) different from that (those) on the *y* axis, because the points in Fig 2 do not show a strong linear relationship. However, Fig 2 shows that *Pa* can be clearly detected by infrared light with specific wavenumbers passing through the surrounding air.

GCMS has been used to show that methyl thiocyanate and benzonitrile are characteristic VMs of *Pa* [20]. These VMs produce strong infrared absorption peaks in the region of 2000–2300 cm^-1^, where one *Pa* index (P at 2213.2 – B at 2213.7) appears. Typical VMs, CO, and N_2_O also give strong infrared absorption peaks in the same region. However, this study does not exclude other VMs that may produce the index peak, because FTIR does not require the ionization of VMs and can provide information about unstable VMs, unlike GCMS.

### Acinetobacter baumannii (Ab)

Fig 3 shows the result for discrimination of *Ab* using two intensities as the *Ab* indexes. One is the peak intensity at 1215.0 cm^-1^ which the subtracted value P at 1215.0 – B at 1214.5, as shown on the horizontal (*x*) axis, and the other is P at 2982.3 – B at 2982.9, as shown on the vertical (*y*) axis. The points represent gas samples including the VMs from *Ab* (filled grey square mark 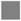) that are shown in the region between the two dotted lines *(y* > 3.76*x* and *y* < 9.65*x*). Almost all the other cases are distinguished.

**Fig 3.**
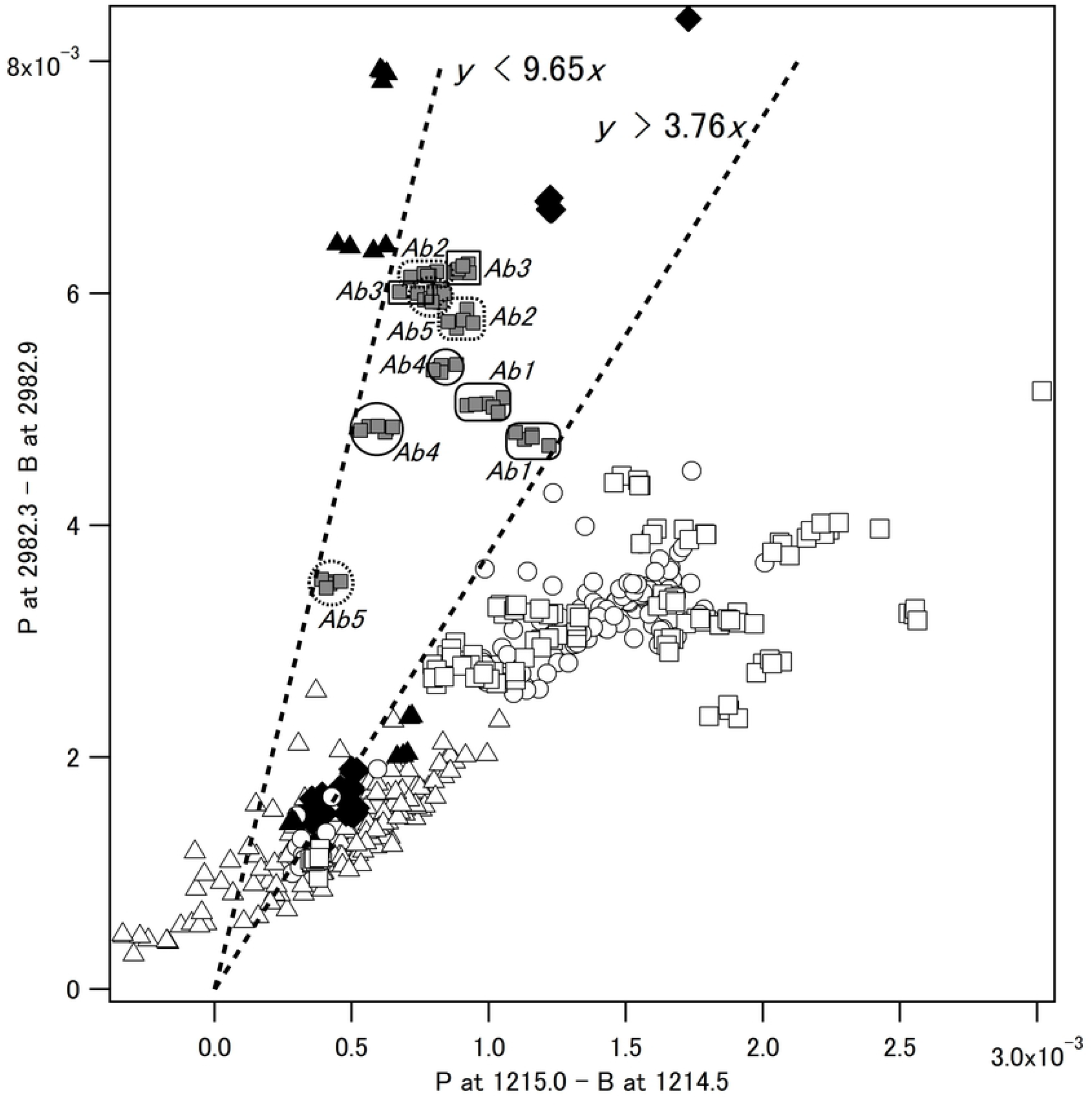
Discrimination of *Acinetobacter baumannii (Ab)* with two *Ab* indexes, P at 1215.0 - B at 1214.5 and P at 2982.3 - B at 2982.9, shown on the horizontal (*x*) and vertical (*y*) axes, respectively. Grey squares 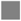, the values obtained from the gas samples that included VMs from *Acinetobacter baumannii (A/b*; black triangles ▲, air from the vicinity of the culture medium alone; black diamonds ♦, *Escherichia coli(Ed;* open squares □, the other 6 types of bacteria in Table 1 except *Ab* and *Ec,* open triangles Δ, the normal room air; and open circles 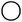, human breath.

The *Ab* index, the intensity of the absorption peak at 2982.3 cm^-1^ is in the CH stretching vibration region; this indicates that the causative VM molecule contains a hydrocarbon structure characteristic of *Ab*. The other *Ab* index observed at 1215.0 cm^-1^ is derived from another VM and the ratio of these VMs is specific to *Ab*, as shown in Fig 3.

In Fig 3, the *Ab* area (*y* > 3.76*x* and *y* < 9.65*x*, a triangular zone between the two dotted lines) overlaps with other plots in the lower left corner. This is due to the detection sensitivity and the wavenumber resolution of the FTIR spectroscopy and it could be improved by adopting an infrared laser system. The *Ab* area also overlaps with the areas for *Escherichia coli (Ec*, ♦ filled black diamond) and the blood agar medium (▲ filled black triangle in the upper left), which causes the *Ab* discrimination rate to deteriorate.

Two additional indices of *Ab* were found, as shown in Fig 4, indicating there was an infrared absorption peak at 4768.7 cm^-1^ (P at 4768.7 – B at 4768.1) on the *x* axis and 5353.8 cm^-1^ (P at 5353.8 – B at 5354.4) on the *y* axis. The two dotted lines (*y* = −188*x* and *y* = 78*x*) separate the points for *Ab*, *Ec*, and culture medium only. By combining the information from Figs 3 and 4, the rate of *Ab* discrimination can be increased.

**Fig 4.**
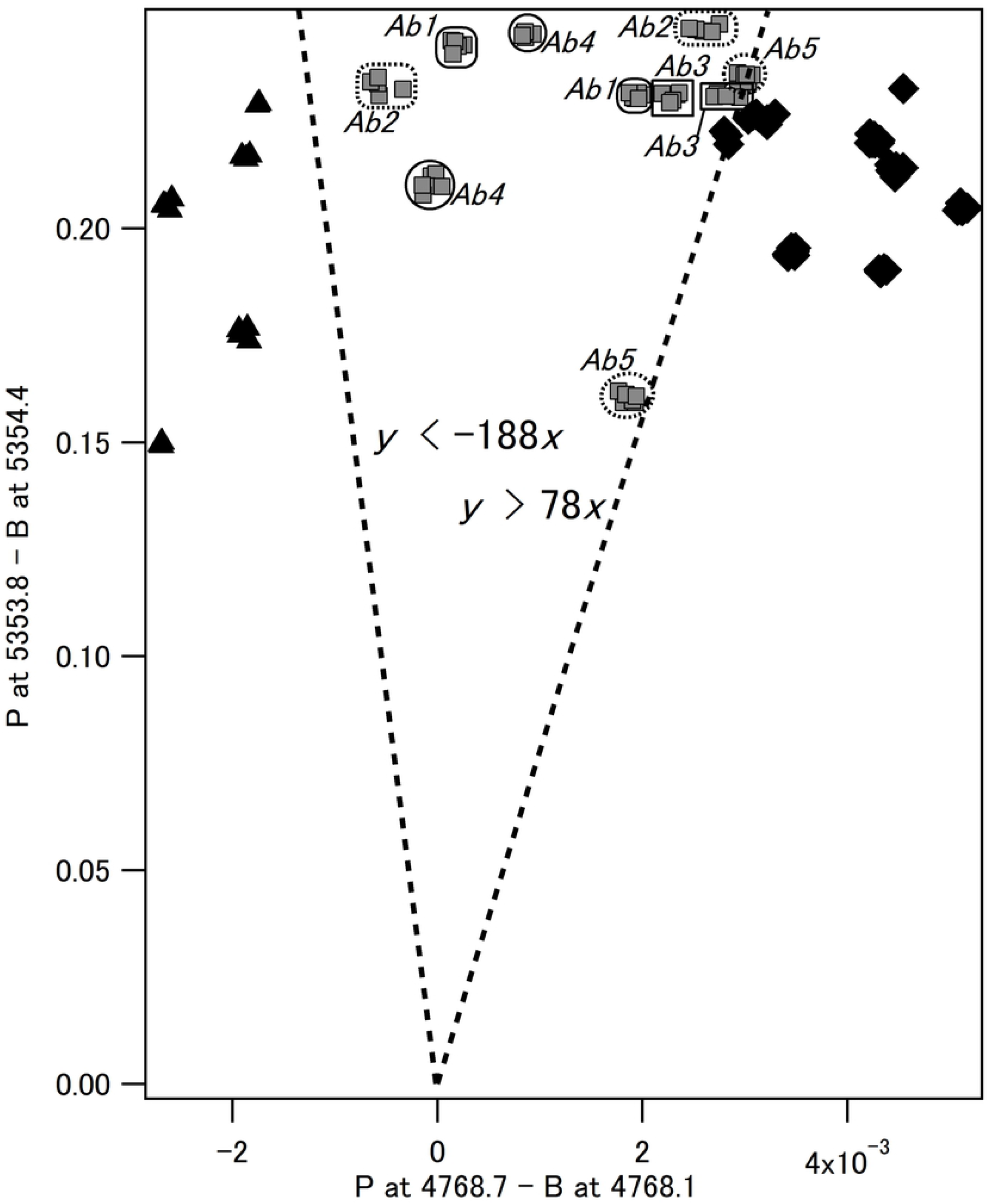
Discrimination between *Acinetobacter baumannii (Ab), Escherichia coli (Ec),* and the blood agar medium only, with two additional indexes P at 4768.7 - B at 4768.1 and P at 5353.8 - B at 5354.4, shown on the horizontal (*x*) and vertical (*y*) axes, respectively. Grey squares 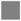, the values obtained from IR spectra for the gas samples including the VMs from *Acinetobacter baumannii (Ab);* black triangles ▲, air from the vicinity of the culture medium alone; and black diamonds ♦, *Escherichia coli (Ed).*

In Fig 4, one of the dotted straight lines has a negative slope, and some plots are in the negative region of the horizontal axis (*x*). Typically, the value of P at 4768.7 – B at 4768.1 is considered positive, because the wavenumber positions at 4768.7 and 4768.1 are assumed to be at the peak (P) and bottom (B), respectively. In fact, none of the gas samples gave the expected infrared spectra. Some gas samples contain a VM that gives an infrared absorption peak where we assume the bottom. This is a reason why not all discrimination lines are ideally based on Lambert-Beer’s law.

## Application and data processing

Infrared lasers can be applied to detect the key VMs of *Pa* and *Ab* using three steps:

1. fine adjustments of the oscillation wavenumbers at the absorption peak (P) and base point (B);
2. conversions of the outputs from the light receiving parts in the infrared laser system to values comparable to the results in this report; and
3. a discrimination program with consideration of the optical path length.

It is worth noting that the gas samples in this experiment were under atmospheric pressure. This work assumes that the atmospheric pressure during any application is the same as at the experiment site.

To replace the absorbance (*A*) obtained by FTIR spectroscopy with the intensity (*I*) obtained from the infrared laser system, the wavenumber positions of P and B were reported as a pair. The relationship between the absorbance (*A*) and transmittance (*T*) is

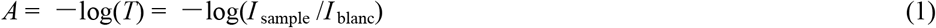

where *I*_sample_ and *I*_blanc_ are the raw values of intensity with and without a sample, respectively. In addition,

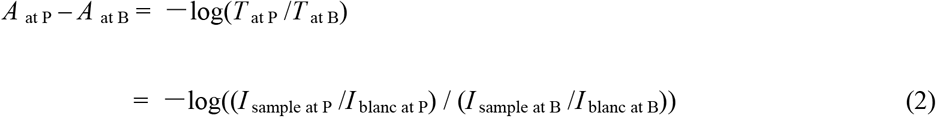

where *A*_at P_ and *A*_at B_ are the absorption values at P and B, respectively.

Infrared lasers have a narrow linewidth and do not give the intensity other than a specific wavenumber. In contrast, FTIR spectroscopy produces a continuous spectrum and the intensity of the infrared light source changes slightly with the wavenumber. In the absence of absorption effects, within a 1 cm^-1^ range, the intensity is assumed to be constant; that is, *I*_blanc at P_ ≈ *I*_blanc at B_. Therefore, from (2) it is possible to obtain

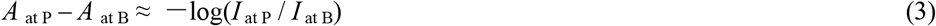

where *I* _at P_ and *I* _at B_ are the raw intensity values at P and B, respectively, which can be observed by the infrared laser system.

The value of *A* _at P_ – *A* _at B_ reflects to the partial pressure (concentration) of a key VM. Two sets of *A* _at P_ and *A* _at B_ for two key VMs can provide information about their component ratios. An appropriate combination of these sets can be used to show the characteristics of the gas sample more clearly.

The discriminant in Figs 2–4 is based on the measurements obtained using a gas cell with an optical path length of 10 m. According to Lambert-Beer’s law, the absorbance is proportional to the optical path length under the same pressure. If the optical path length is L m, which is part of the discriminant, then *y* > 0.003 shown in Fig 2 becomes *y* > 0.003×L / 10. In addition, *x* > 0 in Fig 2, *y* = 9.65*x* and *y* = 3.76*x* in Fig 3, and *y* = −188*x* and *y* = 78*x* in Fig 4 are linear functions through the origin that are not dependent on the optical length.

Since the absorption intensity is proportional to the optical path length, the longer the optical path length, the better the sensitivity. For this reason, infrared laser systems have an advantage over other methods for analyzing low-density gas samples. Most biogas samples are obtained non-invasively. Further, infrared light generally has low energy and is non-destructive. Therefore, it would be realistic to monitor the environment in an open-path space by passing infrared laser light in medical, clinical, and general life settings. When optimizing the optical path according to the required sensitivity and the size of the space to be examined, multiple reflections can be used to increase the optical path length even in a small space, and to achieve high sensitivity.

## Conclusion

This study demonstrated that *P. aeruginosa* and *A. baumannii* can be detected using infrared lights with specific wavenumbers that pass through the surrounding air. The discrimination rates can be improved by observing additional infrared absorptions caused by different key VMs. The results can be used in the application of infrared lasers for continuous monitoring of *P. aeruginosa* and *A. baumannii* with high sensitivity.

The next step involves fine-tuning of the oscillating wavenumber of the infrared laser around the values reported in this study. The fine adjustment will be performed by finding a wavenumber position with the best discrimination rate in consideration of the line width of the infrared laser. As the result, comparisons with traditional methods will be possible.

Applications for human health care will require the accumulation of more detailed information in the infrared spectrum for gas samples under more diverse conditions. We believe that information related to human health, which is statistically significant and has a high discrimination rate, can be extracted by enhancement of the database. At present, we are working with hospitals to accumulate infrared spectra of biogas samples from patients.

## Acknowledgments

We would like to thank Editage for English language editing. This work was supported by JSPS KAKENHI (Grant Number JP26462957) and Research Fund of Showa University, College of Arts and Sciences (2015).

